# Assortative mating in hybrid zones is remarkably ineffective in promoting speciation

**DOI:** 10.1101/637678

**Authors:** Darren E. Irwin

## Abstract

Assortative mating and other forms of partial prezygotic isolation are often viewed as being more important than partial postzygotic isolation (low fitness of hybrids) early in the process of speciation. Here I simulate secondary contact between two populations (‘species’) to examine effects of pre- and postzygotic isolation in preventing blending. A small reduction in hybrid fitness (e.g., 10%) produces a narrower hybrid zone than a strong but imperfect mating preference (e.g., 10x stronger preference for conspecific over heterospecific mates). This is because, in the latter case, rare F1 hybrids find each other attractive (due to assortative mating), leading to the gradual buildup of a full continuum of intermediates between the two species. The cline is narrower than would result from purely neutral diffusion over the same number of generations, largely due to the frequency-dependent mating disadvantage of individuals of rare mating types. Hybrids tend to pay this cost of rarity more than pure individuals, meaning there is an induced postzygotic isolation effect of assortative mating. These results prompt a questioning of the concept of partial prezygotic isolation, since it is not very isolating unless there is also postzygotic isolation.

## Introduction

Speciation, or the evolution of multiple species from one, is usually considered to proceed via the evolution of reproductive isolation between populations (Dobzhansky 1940; Mayr 1942; Coyne and Orr 2004; Rieseberg and Willis 2007; Price 2008). Types of reproductive isolation have traditionally been classified into two main categories. These are (1) prezygotic isolating barriers, which prevent the formation of hybrids, and (2) postzygotic isolating barriers, which cause lower fitness of hybrids compared to members of the parental populations. This categorization is due in part to the idea that they differ in the way selection can act on them: there cannot be direct selection for low hybrid fitness (except possibly in species with substantial parental care; Coyne 1974), whereas there can be direct selection for prezygotic isolation when hybrid fitness is low (Dobzhansky 1940; Mayr 1942; Howard 1993; Liou and Price 1994; Servedio 2000; Hopkins 2013). This latter idea is the theory of reinforcement, which has been controversial (Butlin 1987; Rice and Hostert 1993; Kirkpatrick and Servedio 1999; Servedio and Noor 2003; Servedio 2004) despite providing the logical basis for categorizing isolating factors as pre- or postzygotic.

In much of the literature on speciation, prezygotic isolation is often thought to play a more important role than postzygotic isolation in the initial stages of speciation (Mayr 1942, 1963; West-Eberhard 1983; Grant and Grant 1997; Edwards et al. 2005; Rieseberg and Willis 2007; Schumer et al. 2017; Gill et al. 2019). For example, Mayr (1963) wrote that “ethological barriers to random mating constitute the largest and most important class of isolating mechanisms in animals.” Grant and Grant (1997) stated that “speciation in birds proceeds with the evolution of behavioral barriers to interbreeding; postmating isolation usually evolves much later.” Irwin (2000) suggested that “divergence in mating signals occurs rapidly and can quickly generate reproductive isolation.” Schumer et al. (2017) stated that “premating isolation can play an important role in the early stages of speciation, allowing for the accumulation of other isolating mechanisms.” The widely held idea that assortative mating is important early in speciation appears to result from two patterns in nature: First, traits involved in mate choice and other social dynamics are often the most noticeably divergent traits between populations (e.g., Jones 1997; Irwin et al. 2001; Masta and Maddison 2002). Second, studies quantifying hybrid inviability and infertility have often implied that these postzygotic barriers take a very long time to evolve, and after substantial prezygotic isolation has developed (Coyne and Orr 1989; Price and Bouvier 2002).

These patterns have suggested a key role for sexual signaling and sexual selection in speciation, through their potential to enhance assortative mating. Some theory and empirical patterns support this idea (West-Eberhard 1983; Panhuis et al. 2001; Ritchie 2007; Seddon et al. 2013), whereas others do not (Huang and Rabosky 2014; Servedio and Burger 2014; Cooney et al. 2017). While it is also possible that sexual selection and differentiation in sexual signals impacts fitness of hybrids (Mayr 1942; Kirkpatrick and Servedio 1999; Kawata and Yoshimura 2000; Servedio and Noor 2003; Bridle et al. 2006; Price 2008), this has been less emphasized compared to the role of sexual selection in leading to assortative mating.

Sister taxa tend to be geographically structured, either in complete allopatry or in parapatry, providing strong evidence for a role for geographic separation in speciation (Jordan 1905; Mayr 1942; Price 2008). Range expansion and secondary contact between related populations is common, often leading to the formation of hybrid zones. Such hybrid zones are used as “natural laboratories” for the study of speciation (Hewitt 1988), as they allow the observation of whether the groups are blending and what factors maintain discrete species in the face of ongoing gene flow.

Cline theory has been developed as a tool for describing and understanding hybrid zones (Haldane 1948; Bazykin 1969; Barton 1979, 1983; Barton and Hewitt 1989; Barton and Gale 1993). A sigmoidal cline function can be well fit to many empirical zones, and the width of the zone as well as patterns of linkage disequilibrium can be used to infer whether there is selection maintaining the narrowness of hybrid zones, or conversely whether a zone is neutrally blending through diffusion. One key result is that in a hybrid zone under equilibrium and with random mating within localities, the expected width (*w*) of the zone is (inversely) related to the square root of the strength of selection against hybrids (Bazykin 1969; Barton 1983; Barton and Hewitt 1989):

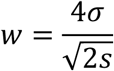

where *σ* is the root-mean-square dispersal distance and *s* is the effective selection against heterozygotes at a locus that is initially fixed for different alleles in the two populations. This type of zone is called a “tension zone,” in which the narrowness is maintained by selection against hybrids balanced by dispersal of parental individuals into the zone (Barton and Hewitt 1989).

Despite the widespread use of cline theory to infer selection in hybrid zones (e.g., Moore and Buchanan 1985; Hewitt 1988; Mallet et al. 1990; Alexandrino et al. 2005; Gay et al. 2008; Brelsford and Irwin 2009; Grossen et al. 2016), it requires a number of assumptions that may be questionable in specific empirical systems. One of these is the assumption of random mating at each location along the cline, ironic given the widespread opinion (see above) that prezygotic isolation is so important in the early stages of speciation. Prezygotic isolation might result from differences in mating signals and preferences, or from differences in ecological traits that lead to habitat-based or timing-based assortative mating.

Here, I use simulations to understand how assortative mating affects the amount of blending between two differentiated populations that come into secondary contact. Do mating preferences for genetically similar individuals results in a narrower cline (compared to the neutral case of random mating), as expected of something that contributes to speciation? I compare the isolating impact of assortative mating to the isolating impact of low hybrid fitness, and examine whether there is a synergistic effect between assortative mating and low hybrid fitness on total isolation of the two populations. The assortative mating modeled here could be seen as being based on a wide variety of underlying mechanisms, for example an active cognitive choice, a physiological mechanism that results in sperm or pollen precedence, or mating within small-scale microhabitats or at specific times that result in similar genotypes tending to mate with each other.

## Methods

The purpose of the simulations presented here is to compare the effects of (1) assortative mating and (2) low hybrid fitness, as well as their combination, on the amount of blending between two species that are coming into secondary contact. The two species are assumed to be ecologically equivalent and the large-scale environment is assumed to be constant in space and time, such that the effects of assortative mating and/or low hybrid fitness can be examined without effects of ecological differentiation. Note that for convenience I use the term “species” for these two populations coming into contact, but in the neutral case there is not any reproductive isolation; the two groups coming into contact could equally well be called “populations” or “subspecies” as the reader prefers. The simulations are individual-based, such that they have limited population sizes and demographic stochasticity plays a role in their behavior. A custom script (named “HZAM”, for Hybrid Zone with Assortative Mating) was written in R (R Core Team 2014) to run the simulations and graph the results.

Simulations take place on a single geographic dimension of length 1. At the beginning of each simulation, individuals of species A have random locations (uniform distribution) in continuous space between locations 0 and 0.48, and individuals of species B have locations between 0.52 and 1; hence the simulations begin at the time just before two isolated populations come into secondary contact. The full geographic space (between locations 0 and 1) has carrying capacity of *K* individuals (in total between the two species, due to their ecological equivalence; with values of *K* ranging from 2,000 to 16,000 depending on the simulation). Given the limited range of each species, this means that simulations start with 0.48*K* individuals within each species. Individuals are either female or male (with equal numbers of the two sexes at the start of the simulation).

Individuals are diploid, each with multiple loci that follow rules of Mendelian inheritance and are not physically linked. Each locus has two alleles, designated 0 and 1, and there is no mutation. At the beginning of each simulation, species A is fixed for allele 0 and species B is fixed for allele 1, at all loci.

The simulations proceed with cycles of mating, reproduction, survival to adulthood, and dispersal to a breeding location. Generations are non-overlapping.

### THE “NEUTRAL” HYBRID ZONE MODEL

In the neutral case, in which there is no assortative mating and no reduced fitness of hybrids, simulations follow these rules:

#### Mating

Each adult female mates with the male that is geographically closest to her.

#### Reproduction

The number of offspring (or children; *C*) of each female is drawn from a Poisson distribution with a mean 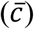 adjusted to reflect local density dependence, a result of intraspecific competition for resources. This is necessary to ensure the population stays spread out in space, and is intuitively appealing because most species are subject to such intraspecific competition (Connell 1983). Local density at a female’s location, *x*_0_, is measured by weighing the locations of all other individuals, *x_real_*, according to a Gaussian curve centered at *x*_0_ with standard deviation

*σ_comp_*:

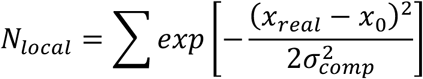

Expected density (i.e., local carrying capacity), *K_local_*, at each location is calculated in a similar way, but assuming *K* individuals are evenly distributed across the range, with locations *x_even_*:

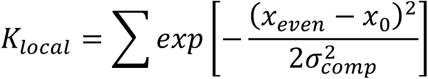

Calculating both real local density and expected local density in this way effectively accounts for edge effects. These expected and actual local densities were then used to calculate the expected number of children for the focal female 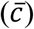 according to the discrete time analog of the continuous logistic growth equation (Prout 1978; Liou and Price 1994):

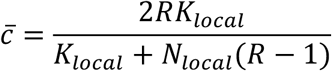

where *R* is the growth rate when the population is small. The 2 in the numerator is due to the fact that only females have offspring, yet they produce both females and males. In the simulations presented here, *R* = 1.05 (i.e., an expected 5% per generation growth rate when the population density is low) and *σ_comp_* = 0.01. Sex of each offspring is determined randomly, with a 50% probability of each.

#### Dispersal

Each offspring acquires an adult breeding location determined by a random draw from a normal distribution centered on the mother’s breeding location, with standard deviation *σ_disp_* = 0.01 (i.e., 1% of the full geographic space). Draws that result in locations larger than 1 or smaller than zero are repeated until the location is within the range.

#### Survival

In this basic neutral model, all offspring survive to adulthood.

### ADDING FUNCTIONAL LOCI TO THE MODEL

Expanding from that basic neutral model, we can model two types of functional effects of loci (Fig. 1):

**Figure 1:**
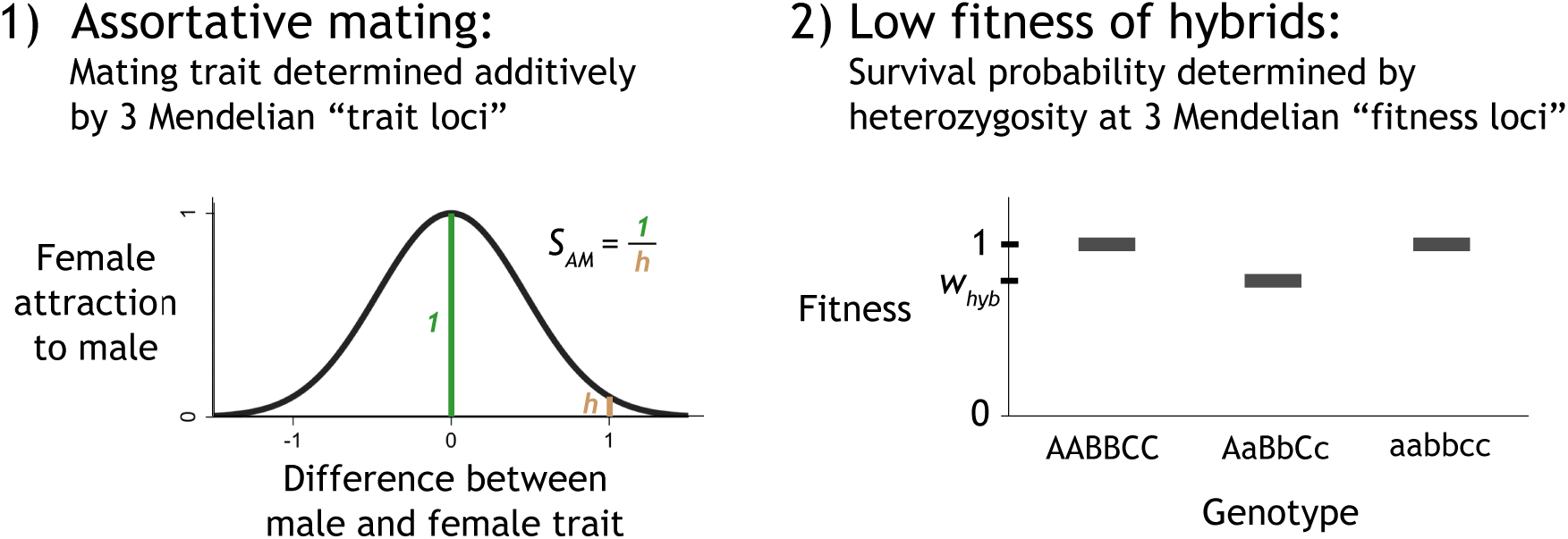
Functional loci in the simulations can cause (1) assortative mating, and/or (2) lower fitness of hybrids. Meanings of the parameters *S_AM_* and *w_hyb_* are illustrated (see also Table 1).

**Table 1:**
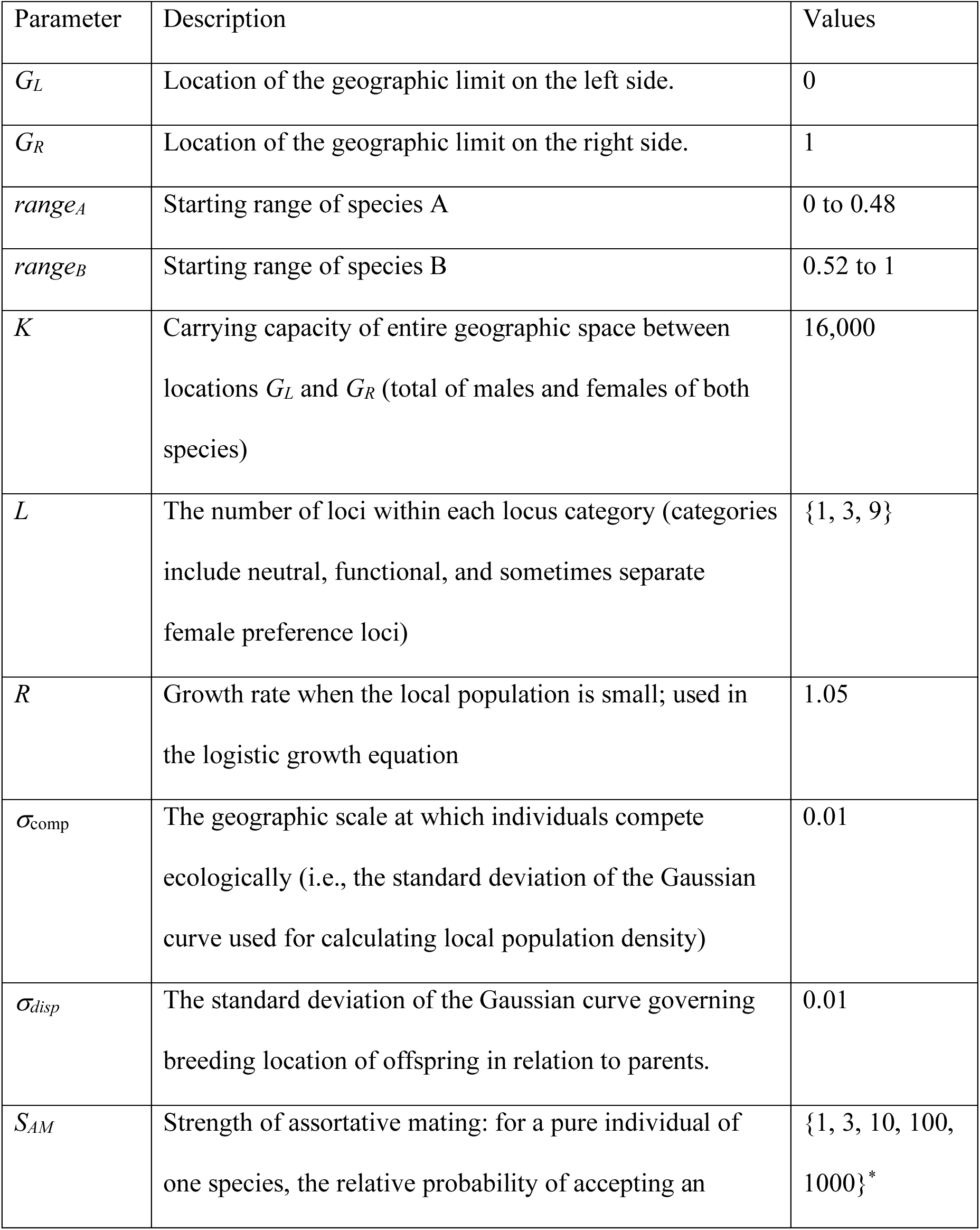
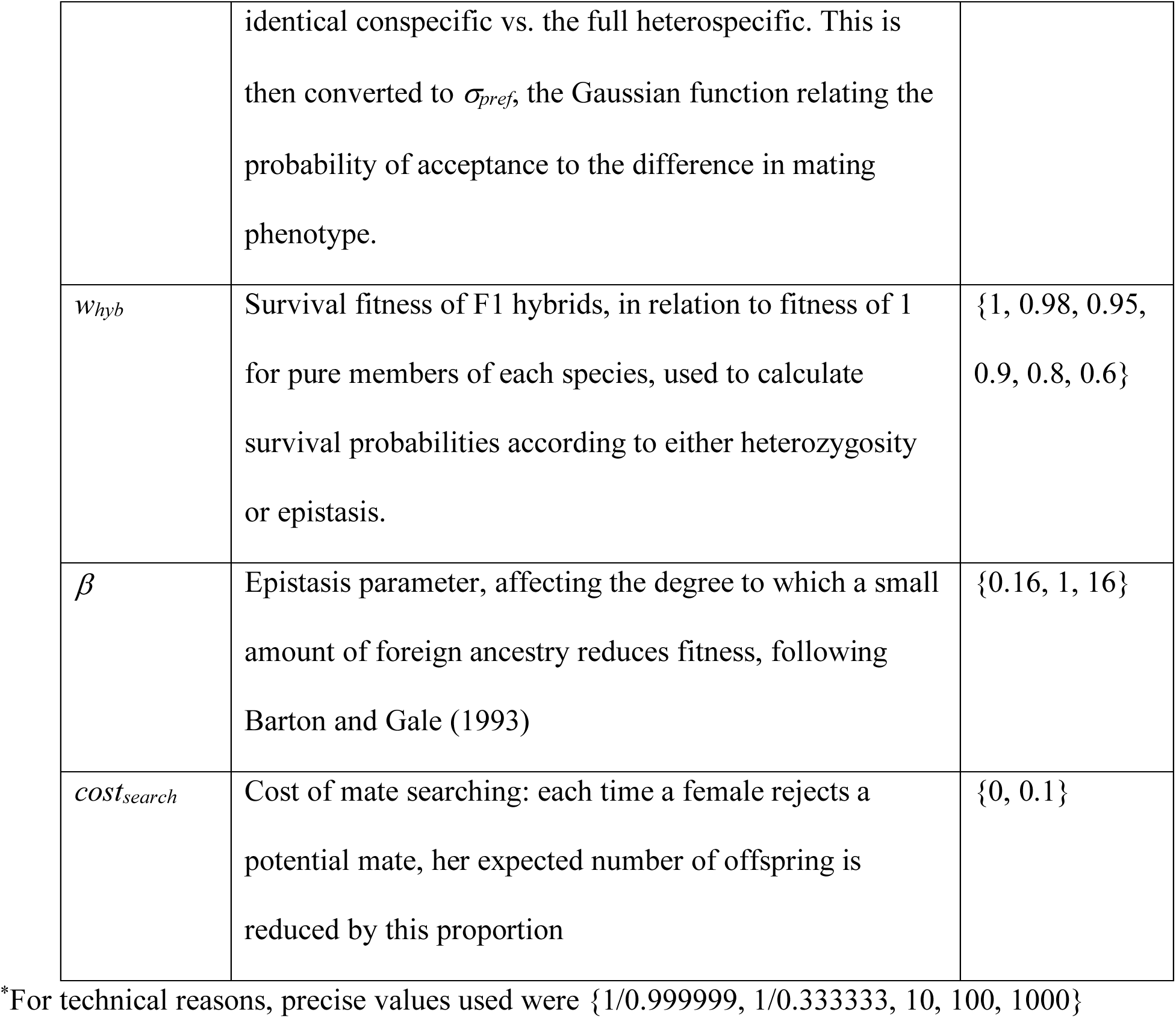
Model parameters and their values as used in the simulations

#### Assortative mating

One or more loci perfectly encode a unidimensional mating phenotype that can have values from 0 to 1. In most simulations presented, alleles affect the phenotype in an additive way, both within and between loci (unless otherwise specified). In a subset of simulations, I consider the effects of dominance by designating alleles from species B (i.e., “1” alleles) as dominant over those from species A (“0” alleles). The mating phenotype can be envisioned as a sexual signal and preference for that signal, or a trait related to timing or microhabitat of breeding. To keep with the bulk of the literature, I explain this in terms of females choosing males based on this trait, although results are likely to be similar if males choose females. When a female encounters a potential mate (i.e., the closest male to a female’s breeding location, initially), she compares his mating phenotype to hers. If a perfect match, she accepts him. If there is some phenotypic difference, her probability of accepting him is determined by a Gaussian function centered on her own phenotype with standard deviation *σ_pref_* (specified in the same units as the mating phenotype). If she rejects him, she then encounters the next-closest male, and repeats this procedure of comparing phenotypes and determining if he is accepted. This continues until she accepts a male. These rules mean that every female finds a mating partner, whereas there is variation in the number of mating partners among males (with some not mating at all). In setting up the simulations and presenting results, I specify the strength of assortative mating (*S_AM_*) in terms of the ratio of the probability of accepting an identical individual as a mate over the probability of accepting a full heterospecific (that is, an individual that is fully 1 unit of mating phenotype away from the choosing individual; see Fig. 1). Most simulations presented have a single mating phenotype that is common to both males and females (and encoded by the same loci), but I also explore the effects of separately encoded male traits and female preferences.

#### Low hybrid fitness

Reduced fitness of hybrids is modeled as reduced probability of survival compared to the pure forms, which have probability of survival equal to one. This lower survival fitness is determined as a result of either underdominance (heterozygotes at each locus have lower probability of survival to adulthood than pure homozygotes) or epistasis (which includes between-gene incompatibilities between alleles from different source populations). In each case, the fitness of F1 hybrids between the two pure starting populations is specified as *w_hyb_*, whereas members of the pure populations have probability of survival equal to 1 (see Fig. 1). In the underdominance case, the loss in fitness due to each heterozygous locus is determined as *s_locus_* = 1 - *w_hyb_*^(1/*U*)^, where *U* is the number of underdominant loci. This enables the calculation of survival probability for individuals heterozygous in only a fraction of loci (e.g., F2’s and backcrosses). Thus the probability of each offspring living to adulthood is determined as *p_surv_* = (1 - *s_locus_*)*^H^*, where *H* is the number of heterozygous underdominant loci in that individual. In the epistasis case, survival fitness is modelled following Barton and Gale (1993) as *p_surv_* = 1 - (1 - *w_hyb_*)(4*x*[1 - *x*])*^β^*, where x is the fraction of alleles derived from one of the populations and *β* is a parameter that influences the degree to which a small amount of foreign ancestry reduces fitness (I focus on the simplest case, with *β* = 1, and also show results for *β* = 16 and *β* = 1/16).

In the simulations presented here, functional alleles can have effects on either assortative mating or low hybrid fitness, or on both. In the latter case, the same loci have both types of functional effects (rather than having different loci affect each). In the main set of simulations presented, there are 3 functional loci and 3 neutral loci, all not physically linked (i.e., recombining freely).

### CLINE FITTING AND WIDTH QUANTIFICATION

During and after the simulations, the shape of the hybrid zone is estimated by fitting a generalized additive model (GAM) to the relationship between geographic location and the “hybrid index” (*HI*) value of individuals, defined as the average value of alleles within each individual. Fitting is done using the “ML” method in the “gam” command of the R package “mgcv” with a quasibinomial error structure and logit link.

To quantify width of the hybrid zone, I define the hybrid zone to include the geographic area between the locations at which the GAM cline fit crosses *HI* = 0.1 and *HI* = 0.9. This somewhat arbitrary definition was chosen because it produces values that are reasonably similar to other definitions of hybrid zone width in the literature (e.g., the inverse of the steepest slope; Barton and Hewitt 1985) but appears less subject to stochastic noise. All comparisons in this paper are between simulations run using this method of quantifying width.

There are two ways the width of a contact zone, as measured above, can become large. First, an extensive hybrid zone can develop, such that the center of the zone contains a range of intermediate genotypes, with few if any pure parentals. Second, under some conditions there can be extensive overlap between two distinct species, with no or few hybrids present, and a gradual change in frequencies of the two species over space. The former can be referred to as a “unimodal” hybrid zone, and the latter as a “bimodal” overlap zone (or hybrid zone if some hybrids are observed; Jiggins and Mallet 2000; Harrison and Bogdanowicz 2006). To distinguish these, at the end of each simulation I record the numbers and characteristics of individuals in the center of the zone. For this purpose, the center is defined as the region within half a dispersal distance (i.e., 0.5*σ_disp_* = 0.005) from the location at which the cline fit crosses *HI* = 0.5. I then calculate bimodality as the proportion of individuals in the center who have pure genotypes (that is, individuals with either all 0 alleles or all 1 alleles).

In most simulations presented, there are two categories of loci: functional loci that can have direct effects on assortative mating and/or fitness of hybrids; and neutral loci that have no functional effects. Clines can be fit to each category of loci, such that widths of clines between functional and neutral loci can be compared. Cline widths are recorded once every five generations during each simulation, enabling the time course of hybrid zone expansion to be recorded.

Model parameters and the values used in the simulations are summarized in Table 1. In each of the main modeled scenarios, I present results from five replicate simulations, each lasting 250 generations.

## Results

I start by examining and comparing two scenarios: (1) moderately strong assortative mating, such that a female of species A is 10 times more likely to accept a male of species A than a male of species B (this is called “10x assortative mating”), and (2) modest selection against full heterozygotes (i.e., F1 hybrids, or F2’s or other individuals that are fully heterozygous at the underdominant loci) of 10% (called “90% hybrid fitness”).

Under 10x assortative mating, hybridization is very rare in the first few generations of contact. However, rare hybridization leads to some F1 individuals forming in the contact zone between the two species. Given the rules of assortative mating (additive, with Mendelian inheritance of a single phenotype that determines male trait and female preference), the F1 individuals often pair with other F1 individuals, leading to F2’s. Moreover, F1’s are more attractive to each parental species than the two species are to each other, leading to backcrosses. These dynamics continue and gradually lead to the buildup of all combinations of intermediate forms, a “genetic bridge” between the species (Fig. 2, left panels). After the broad hybrid zone develops, individuals from the pure species rarely come into contact, as the hybrid zone tends to keep them spatially separated. Individuals with similar HI values find each other to be acceptable mates, such that there is a broad zone of transition between the two species, with intermediates (*HI* = 0.5) common in the middle of the zone.

**Figure 2:**
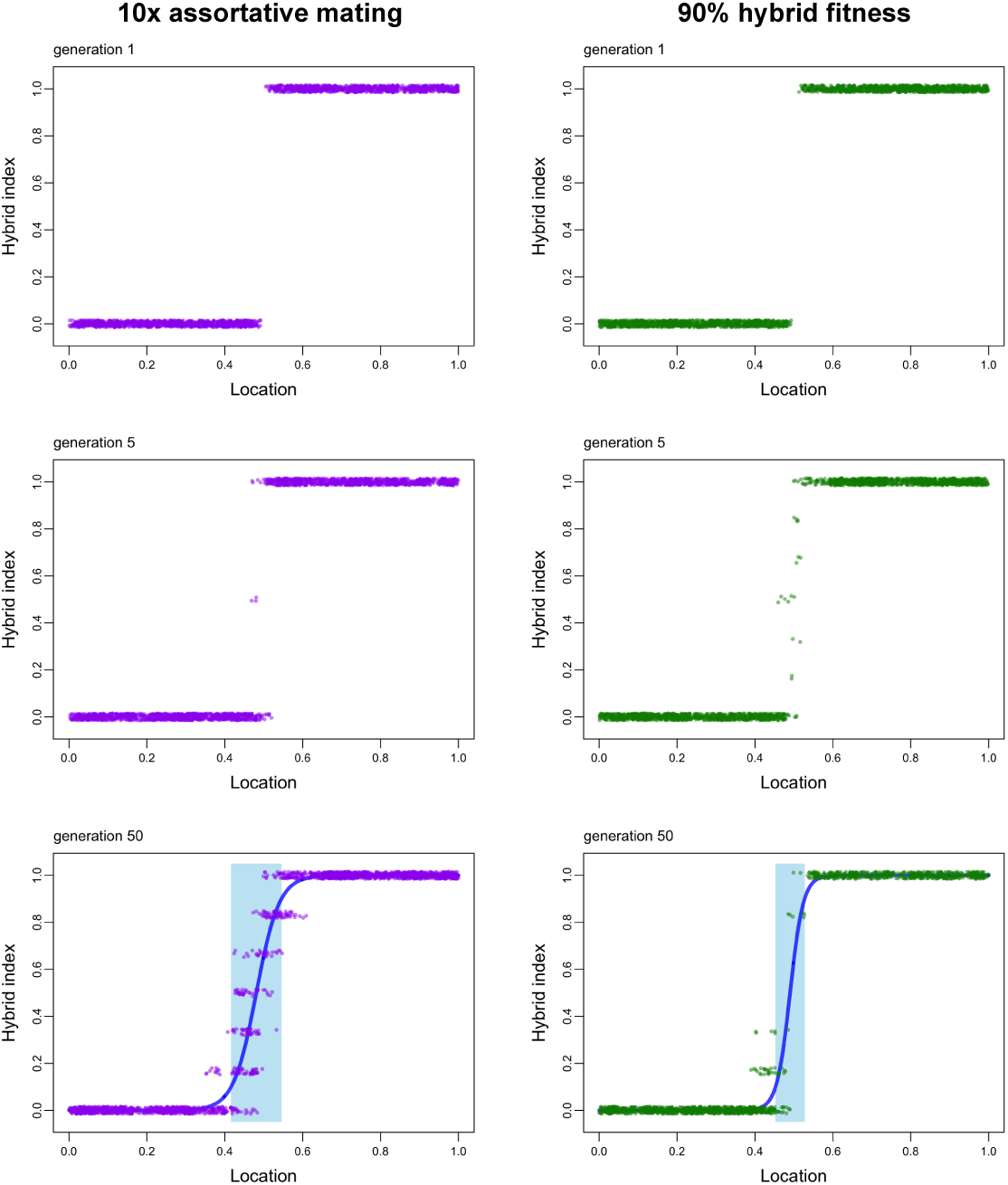
Hybrid zone development under two scenarios: on left, 10x assortative mating (each purple dot represents one individual, with a bit of jitter added to their hybrid index values to show them more clearly); on right, 90% hybrid fitness (each green dot represents one individual). Upper panels show the simulation at generation 1, middle panels at generation 5, and lower panels at generation 50. In the latter, the grey area indicates the hybrid zone, as defined by the region in which the cline fit (in blue) has values between *HI* = 0.1 and *HI* = 0.9. Assortative mating causes a delay in the development of the hybrid zone, but eventually the zone is wider in the 10x assortative mating case than in the 90% hybrid fitness case. In these simulations, there are three functional and carrying capacity of the whole range *K* = 2000 (fewer than the K = 16000 used in other figures, in order to better illustrate in this small graph format).

This situation contrasts with that of a small amount of postmating isolation (i.e., reduced fitness) in hybrids, without any assortative mating. In this case (Fig. 2, right panels), hybridization occurs rapidly (because there is no preference for mates who are similar). However, the zone stays narrow over time, and perfect intermediates (*HI* = 0.5) stay somewhat rare due to their lower fitness. These “tension zone” dynamics have been described well by earlier studies (e.g., Barton and Hewitt 1989); the narrowness of the zone results from a balance between selection against hybrids and net greater gene flow into the hybrid zone than out of the zone.

Running these simulations over longer time periods (500 generations each) and averaging results over multiple replicates (3 for each set of parameters) reveals that a modest amount of selection against hybrids (10%) maintains a narrower hybrid zone that moderately strong (10x) assortative mating (Fig. 3A).

**Figure 3:**
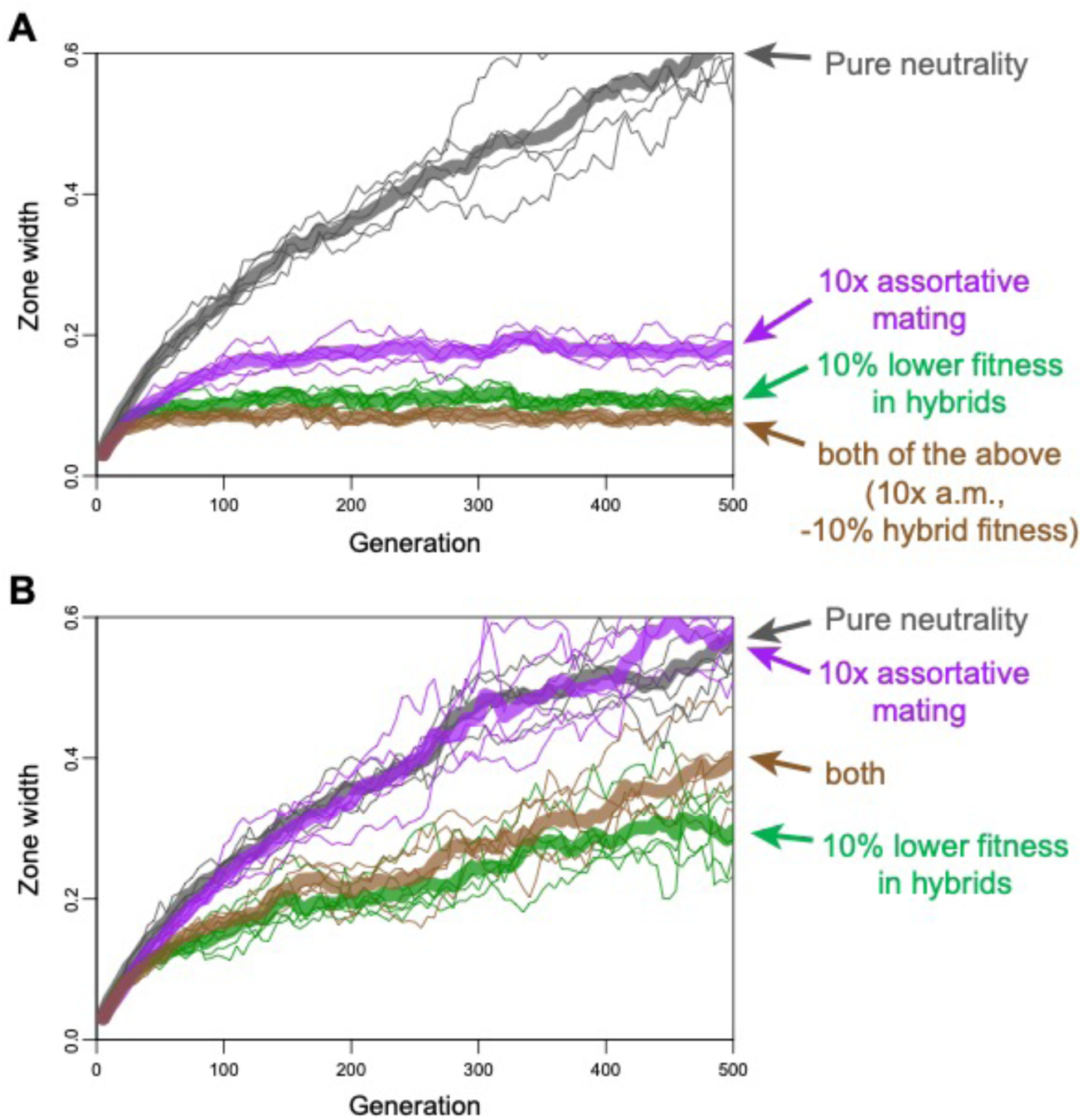
Relationships between hybrid zone width and time (i.e., number of generations) following secondary contact of two species, for (A) functional loci and (B) neutral loci. For each condition, five replicate simulations are shown (narrow lines of the same color) as well as their mean (broad transparent lines). For neutral loci, only conditions involving lower fitness of hybrids noticeably suppress zone width, whereas assortative mating alone has no noticeable effect (compared to the fully neutral model) on zone width. For these simulations, *K* =16000, and there are 3 functional loci with the effects as indicated, as well as 3 neutral loci.

While 10x assortative mating is less powerful than 90% hybrid fitness in keeping a hybrid zone narrow, 10x assortative mating does tend to keep the zone narrower than expected under the purely neutral case (Fig. 3A). Much of this effect is due to induced frequency-dependent selection against rare mating types. Outside of the hybrid zone, where each species is fixed for one or the other mating type, all potential mates view all others as completely acceptable. Inside the zone, where there is variation in mating types, each male is maximally acceptable to only a fraction of females near him. Males of a rare mating type (in proportion to female mating types around them) are less likely to be chosen by a female. Given the steepness of the hybrid zone in the middle of the zone, this rare male disadvantage is likely to affect individuals close to *HI* = 0.5 more than those closer to pure forms. If so, this rare-male-type disadvantage can be considered a form of low hybrid fitness, in other words a form of postzygotic isolation, even though it is induced by assortative mating.

So far, the analysis has focused on the cline resulting from functional loci. Another way to view speciation focuses on the restriction of flow of alleles at neutral loci—those with no effects on assortative mating nor fitness—across hybrid zones maintained by selection on functional loci. Figure 3B shows, for each of the conditions already examined, the broadening over time of the width of the cline in neutral loci (which are not physically linked to functional loci). Of these scenarios, only the explicit selection against heterozygotes results in noticeable reduction in neutral loci gene flow compared to the case of a purely neutral hybrid zone. Figure 4 shows a more comprehensive summary of cline widths and bimodality resulting from combinations of hybrid fitness and assortative mating, for both functional and neutral loci (in this case, 3 of each).

**Figure 4:**
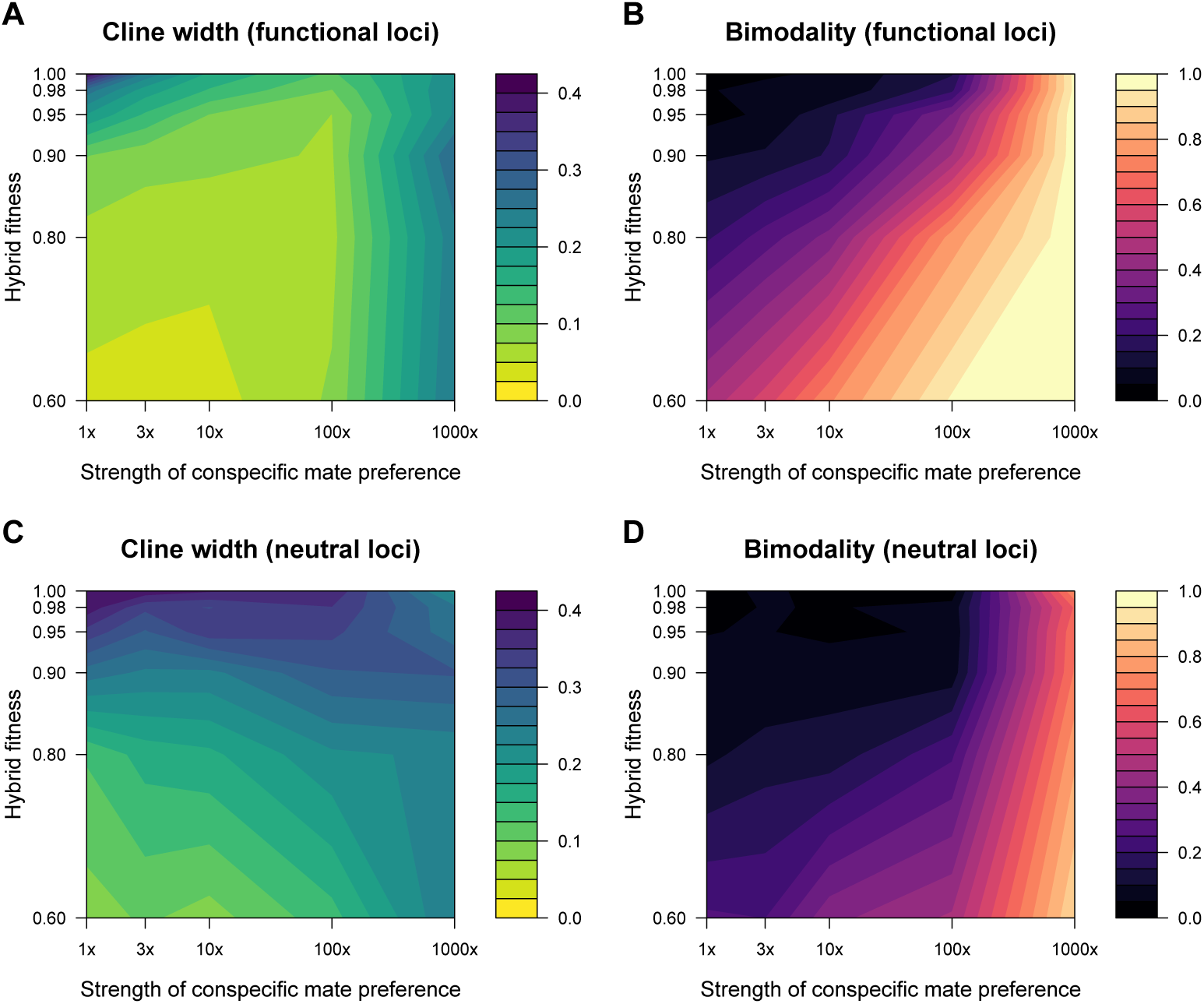
The dependence of hybrid zones characteristics (width and bimodality, measured 250 generations after contact) on the strength of assortative mating and the fitness of hybrids. (A) and (C) show contact zone width, and (B) and (D) show bimodality in the zone center. In (A), we see that a small reduction in hybrid fitness (i.e., down to about 95% that of pure forms) results in much narrowing of the zone, as measured on the functional loci. Assortative mating adds only a little to the narrowing of the zone, except when conspecific mate preference becomes very strong (i.e., about 100 to 1000x), when there is extensive overlap between the pure species. We see this in (B), showing that that zones are unimodal (i.e., mostly intermediates in the center) unless assortative mating is very strong, such that zones become bimodal (mostly pure individuals in the center). In (C), we see that the width of the zone based on neutral loci is narrowed by just a small reduction in hybrid fitness (e.g. from 100% to 95%), but increasing assortative mating strength up to 100x does not result in narrower clines at neutral loci. Only at extreme conspecific mate preference (e.g., 1000x) does the zone become largely bimodal (i.e., an overlap zone with little hybridization), limiting spread of neutral loci between species. These graphs are based on a set of simulations with *K* = 16,000, 3 functional loci (additively encoding mate preference and fitness), 3 neutral loci, and *σ_disp_* = 0.01. Tick marks along the axes indicate the parameters at which combinations were run.

Given the apparent low effectiveness of prezygotic isolation in keeping hybrid zones narrow, both for functional and neutral loci, are there conditions where the prezygotic isolating role of assortative mating has a larger impact? Two regions of parameter space stand out, but both are of questionable relevance to the early stages of speciation.

First, if the assortative mating phenotype (both signal and preference) is encoded by a single locus (rather than three in the above simulations), then the zone width stays much narrower (Fig. 5). However, the relevance of a 1-locus cline to speciation is debatable, as speciation is usually considered to involve differentiation in multiple loci. As the number of loci encoding the assortative mating trait increases, the impact of assortative mating on cline narrowness declines (Figs. 5, S1, S2).

**Figure 5:**
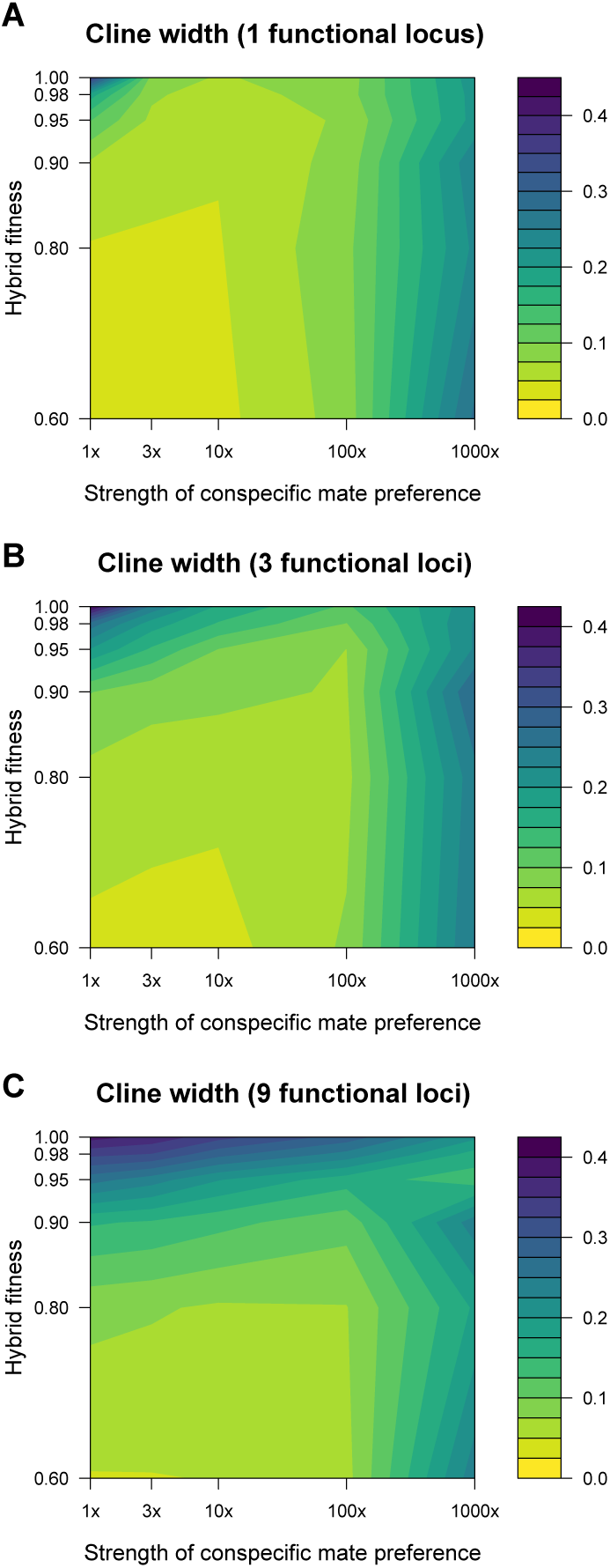
The number of loci encoding a trait influences the relationship between assortative mating, low hybrid fitness, and hybrid zone width. As the number of loci increases from 1 (in A), to 3 (in B), to 9 (in C), there is a decreasing role of conspecific mate preference in limiting hybrid zone width. Aside from the number of loci, all other settings are the same as in Fig. 4. For each response variable, the graphs show the arithmetic mean across five runs for each set of parameters.

Second, if assortative mating is extremely strong, e.g. 100x to 1000x (depending on the strength of selection against hybrids), then hybridization can be so rare that the genetic bridge between species does not form, leaving a bimodal overlap zone (see Fig. 4B). In this case, prezygotic isolation is essentially complete, which makes this scenario somewhat unrelated to the debate over whether prezygotic or postzygotic speciation are more important early in the speciation process, when both are incomplete. Again, however, a major component of the isolation is due to very rare hybrid males not being attractive to females of either species—this can be considered a form of postzygotic isolation.

Factors such as dominance, epistasis, and separate loci for the trait and preference can also be considered in terms of how they affect hybrid zone width and bimodality. In the basic model considered above, F1 hybrids have intermediate mating phenotypes; instead, what if mating traits and preferences of one species are dominant, such that F1 hybrids have a mating phenotype identical to that species? Figures 6A and S3 show the resulting genetic cline widths and bimodality are similar to those in the simple case of codominant loci (compare to Fig. 4). While it might seem that dominance would result in less blending between the species (because F1s are identical phenotypically to one of the parents), dominance can actually result in F1s backcrossing more readily (rather than tending to mate with other hybrids), thereby facilitating gene flow. In the basic model, hybrid survival fitness is determined by heterozygosity; instead, what if there are epistatic interactions between genes? Figure 6B and S4, based on an epistasis parameter (*β*) of 1, again shows a very similar pattern to the basic model (Fig. 4). Extreme values of the epistasis parameter can noticeably affect the impact of low hybrid fitness on cline width, especially for neutral loci (Figs. S5, S6). In the basic model, male traits and female preferences are encoded by the same set of loci; instead, what if there are different loci encoding each? Figures 6C and S7 show that, when there is no reduced fitness of hybrids, there is virtually no impact of mating preferences on keeping a hybrid zone narrow or bimodal, even less than in the basic model (Fig. 4A,B). In the basic model, there are no mate search costs for females; instead, what if female expected fecundity is reduced 10% for each male rejected as a mate? Figures 6D and S8 show this causes mating preferences to have a much larger role in keeping a hybrid zone narrow and bimodal. However, in a hybrid zone this mating search cost tends to be paid more by hybrids than the pure forms, such that it is a form of postzygotic isolation. One interesting consequence of search costs is that when premating isolation is essentially complete (the far right side of each graph in Figure 6D), search costs prevent a wide overlap zone forming, a consequence of frequency-dependent selection due to females of the more rare species paying higher search costs than those of the common species.

**Figure 6:**
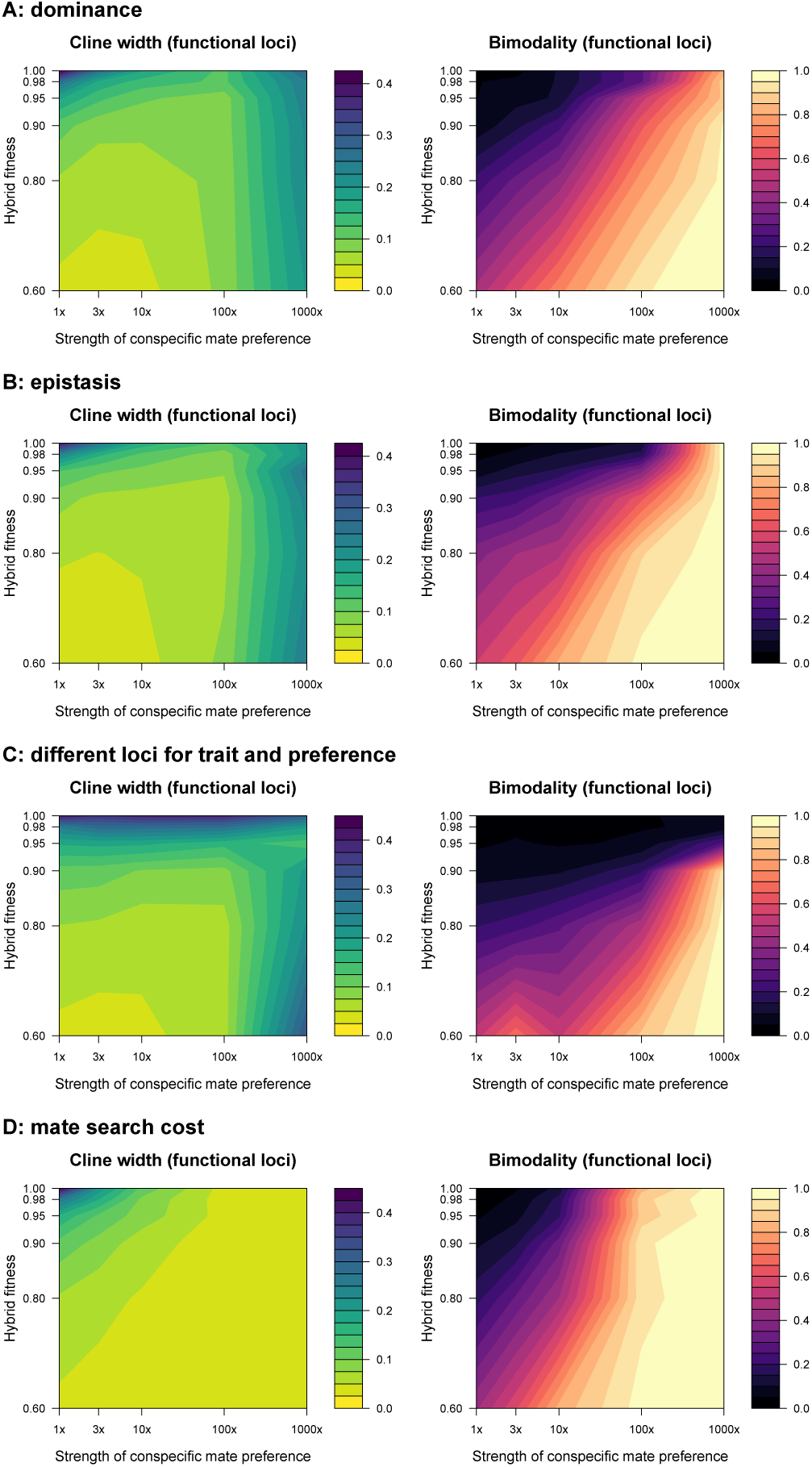
The influence of assortative mating and low hybrid fitness on hybrid zone width and bimodality when there is (A) dominance in the mating phenotype; (B) epistasis (with *β* = 1 rather than simple heterozygote disadvantage in loci determine survival probability; (C) different loci for male trait and female preference (3 loci each), with cline width and bimodality shown for the male trait loci, and the survival fitness based on those same loci; or (D) female mate search cost of 10% reduced reproductive fitness per male rejected. All other settings are the same as in Figure 4. For each response variable, the graphs show the arithmetic mean across five runs for each set of parameters. See figures S3-S8 for more details.

## Discussion

When two somewhat differentiated populations come into contact, their interactions determine whether the two forms blend back together or persist as distinct entities that may continue to diverge into two species. Much of the literature implies there are two main categories of “reproductive isolating mechanisms” that can prevent blending: prezygotic (e.g., assortative mating) and postzygotic (i.e., low hybrid fitness). Previous models of secondary contact involving continuous space and limited dispersal have examined only a subset of possible combinations of assortative mating and low hybrid fitness: tension zone theory has addressed the case of random mating (i.e., zero assortative mating) and partial postzygotic isolation (Bazykin 1969; Barton 1979; Barton and Hewitt 1989; Barton and Gale 1993), whereas Goldberg and Lande (2006) have examined the case of partial assortative mating and complete postzygotic isolation (i.e., 100% inviable hybrids). In both cases, narrow hybrid / contact zones between the populations are maintained due to the low fitness of hybrids (in the case of Goldberg and Lande [2006], a factor limiting range expansion is the larger negative impact of hybridization on the rare species). Here, I have examined a model that can include both partial assortative mating and partial selection again hybrids, allowing inference of the effect of each factor and their combination on the width and bimodality of contact zones. Results indicate that conspecific mate preference, unless essentially complete (meaning a preference of roughly more than 100x to 1000x), has little influence on the suppression of blending between incipient species. In contrast, a small reduction in hybrid fitness has major impact on limiting the width of hybrid zones.

These results can be understood by considering the effects of assortative mating and low hybrid fitness during multiple generations of breeding in a hybrid zone. Reduced hybrid fitness (i.e., partial postzygotic isolation) has an effect that continues through generations following initial hybridization. This results in lower reproductive output in the center of the zone and a resulting net gene flow into the zone. This is the “tension zone” phenomenon well described in the literature (Key 1968; Barton and Hewitt 1985, 1989; Barton and Gale 1993). In contrast, prezygotic isolation (as normally defined) applies only to the interactions between the “pure” species. Assortative mating is usually considered to be a form of prezygotic isolation. Yet if reproductive isolation is incomplete, and if the same rules of assortative mating (i.e., like tends to pair with like) apply to hybrids, then F1 hybrids are attractive to each other. Furthermore, hybrids are more acceptable to each parental species than fully heterospecific individuals are. These dynamics lead to the formation of F2’s, backcrosses, and so forth through the generations until a complete phenotypic and genotypic bridge is formed between the two species. Hence the ability of assortative mating to keep the species genetically separated declines through the generations following initial contact. While the view that prezygotic isolation is more important than postzygotic isolation in the early phases of speciation is common (e.g., Mayr 1942; West-Eberhard 1983; Grant and Grant 1997; Edwards et al. 2005; Rieseberg and Willis 2007; Schumer et al. 2017), Price (2008, p. 399) recognized that “even a low frequency of hybridization should lead to the merging of two species back into one, if hybrids are perfectly fit” (see also Liou and Price 1994).

Several other studies, using widely differing models, have also indicated that strong but incomplete assortative mating can be ineffective in preventing two populations from blending. Singhal and Moritz (2012) used a stepping-stone model of a hybrid zone with assortative mating modeled in the following way: a proportion of individuals (*α*) mate preferentially with individuals sharing at least 90% of their ancestry, and the remaining (1 - *α*) individuals mate randomly. They tested *α* values ranging from 0 to 0.8, and found no significant effect of assortative mating on the widths and characteristics of the hybrid zone. Pulido-Santacruz et al. (2018) used an island model (a single non-spatial hybrid zone, with immigration from two parental islands) to show that even strong assortative mating (up to a factor of a 100x preference for conspecifics) results in extensive blending of two species. Using spatially explicit simulations of hybrid zones involving smaller population sizes than those analyzed here and separate loci encoding male traits and female preferences, Sadedin and Littlejohn (2003) also found little to no effect of the strength of mate choice on the persistence of bimodal hybrid zones.

The model analyzed here is based on continuous space and no geographic variation in environmental variables, as the purpose is to assess the relative impact of assortative mating and low hybrid fitness on the width and bimodality of hybrid zones. Incorporating geographic variation in the environment and local adaptation would introduce a form of indirect postzygotic isolation to the model, as hybrids would often find themselves poorly suited to the local environment. A possible exception is provided by models of ‘bounded hybrid superiority’ hybrid zones (Moore 1977), in which hybrids tend to have high fitness in a band of intermediate habitat between the two parental species; such situations are beyond the scope of the present analysis, as they are not very relevant to the production or maintenance of reproductive isolation (since hybrids thrive in the intermediate zone and provide a conduit for gene flow between the parental species). Habitat heterogeneity is also thought to be important in the formation of ‘mosaic’ hybrid zones, in which there is a complex patchy pattern of the two parental species and hybrids (Harrison and Rand 1989). However, M’Gonigle and Fitzjohn (2010) presented a model in which long-distance dispersal together with assortative mating leads to a mosaic hybrid zone in the absence of ecological differences or intrinsic incompatibilities between species. This in large part is due to rare mating type disadvantage (M’Gonigle and Fitzjohn 2010), which can tend to have a larger effect on hybrids (because they are initially rare) than on the parental species. The magnitude of this effect in the study of M’Gonigle and Fitzjohn (2010) is likely amplified by two aspects of the model: using a demic structure (rather than continuous space) and not allowing hybrids to influence allele frequencies in the parental populations, thereby imposing a limited hybrid zone width.

Given the results presented here, what explains the common perception that prezygotic isolation is more important than postmating isolation early in speciation? For one, sexual signals are often among the most obvious differences between sister species or populations within species. Add to this the tendency of empirical examinations of postzygotic isolation to focus on hybrid inviability and infertility, quite severe forms of reduced hybrid fitness that often take a long time to evolve compared to typical time courses of speciation (Coyne and Orr 1989, 1997; Presgraves 2002; Price and Bouvier 2002; Bolnick and Near 2005; Sasa et al. 2006; Scopece et al. 2008; Turissini et al. 2018). Likewise, many of the theoretical examinations of postzygotic isolation have focused on patterns related to inviability and infertility (e.g., the debates regarding Haldane’s rule and the large-X effect, e.g. Turelli and Orr 1995; Schilthuizen et al. 2011). Because those severe forms of postzygotic isolation take so long to evolve, many have reasoned that postzygotic isolation is not important early in speciation. This would leave prezygotic isolation as the logical cause.

However, there are many forms of partial postzygotic isolation that are far less severe and less noticeable that inviability and infertility. A few examples include physiological problems (due e.g. to cytonuclear discordance; Burton et al. 2013; Hill 2019), reduced cognitive ability (McQuillan et al. 2018), intermediate ecological behaviors (e.g. intermediate seasonal migratory behavior; Delmore and Irwin 2014), intermediate morphology that renders individuals poorly suited to the environment (Benkman 2003; Schluter 2009; Grant and Grant 2014), and problems with reproductive compatibility of hybrids that would be too subtle to be noticed by earlier studies examining very severe infertility (Bridle et al. 2006; Brodin and Haas 2006; Knegt et al. 2017). Servedio and Noor (2003) and Bridle et al. (2006) discussed two distinct forms of ‘behavioral hybrid dysfunction’ (BHD) that can reduce mating success: ‘extrinsic BHD’ results entirely from intermediacy in mating signals and preferences, a result of frequency-dependent disadvantage of rare intermediates (and hence ‘extrinsic’ because based on the social environment), whereas ‘intrinsic BHD’ results from some behavioral deficiency that reduces mating success, beyond simple intermediacy of signal or preference.

A further reason that prezygotic isolation is emphasized as a cause of speciation is the commonly held idea that factors that act earlier in the life cycle are more important. This idea appears to result from the theory of reinforcement: postzygotic isolation causes selection for prezygotic isolation (Dobzhansky 1940; Howard 1993; Liou and Price 1994; Hudson and Price 2014). However, this reasoning necessitates a primary role of postzygotic isolation, rather than prezygotic isolation, in initiating speciation.

These findings prompt a reconsideration of standard approaches to quantifying the relative strengths of different forms of reproductive isolation when both pre- and postzygotic isolation are incomplete. Some commonly used approaches for quantifying prezygotic isolation apply only to a single generation, prior to the buildup of hybrids over generations. For example, the formula given by Coyne and Orr (2004, p. 185) to quantify habitat isolation, a component of prezygotic isolation, is 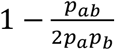, where *p_ab_* is the proportion of heterospecific encounters between individuals of opposite sexes, *p_a_* is the proportion of individuals that are species *a*, and *p_b_* is the proportion of individuals that are species *b*. This formula does not consider the frequency of hybrids, meaning it can only be sensibly applied when hybrids have zero or very low frequency. Formulae for other components of prezygotic isolation (pollinator isolation and temporal isolation) appear on pages 195 and 204 of Coyne and Orr (2004), and likewise leave out hybrids (see Ramsey et al. 2003 and Kay 2006 for similar approaches). The simulations here show that moderately strong levels of prezygotic isolation (e.g., up to 100-1000x preference for conspecific mates) would, in the absence of postzygotic isolation, lead to a complete breakdown of isolation between the two species. Hence the use of these formulae to quantify partial prezygotic isolation is an approach that is only valid when postzygotic isolation is sizeable, which is usually the case in the studies that have used this approach (e.g., Ramsey et al. 2003; Kay 2006).

These considerations raise a number of questions: Can partial prezygotic isolation actually be defined, either conceptually or mathematically, given that it does not prevent extensive blending between species in the absence of postzygotic isolation? What does it mean to say, as Mayr (1963) and others have, that prezygotic barriers are more “important” than postzygotic barriers in keeping species apart? If the goal is simply to examine the breakdown of isolation in a single generation soon after two populations first meet, then it makes sense to use the equations above to quantify the contribution of these barriers to preventing the production of breeding F1 hybrids in the next generation. That approach might reveal that prezygotic isolation is quantitively larger than postzygotic isolation, hence more important during that one generation. In contrast, if the goal is to quantify the contribution of assortative mating and low hybrid fitness to the longer-term maintenance of two distinct species (i.e., important to speciation), then that approach fails. This is because a small reduction in hybrid fitness is highly effective at limiting blending of the species, whereas strong (but incomplete) assortative mating in the absence of postzygotic isolation does not prevent extensive blending of the species. When partial assortative mating does have an effect in preventing blending, it is largely due to induced mating disadvantage of hybrids, a form of postzygotic isolation. Essentially, partial prezygotic isolation is not very isolating, and is unclear how to define or quantify when hybrids are present at appreciable frequencies (i.e., in hybrid zones).

The historical tendency to classify isolating barriers as either prezygotic or postzygotic is based largely on their different relationships with selection. There is not normally direct selection for postzygotic isolation, but there can be for prezygotic isolation if there is low fitness of hybrids (Dobzhansky 1940; Mayr 1942). This is the theory of reinforcement (Dobzhansky 1940; Howard 1993; Hopkins 2013). Some previous analyses (based on island models) concluded that reinforcement can only occur if postzygotic isolation is quite strong (Liou and Price 1994; Servedio 2000). Such a condition is unlikely in the early stages of speciation, but it is increasingly likely in the latter stages. Results from simulations presented here suggest that strong assortative mating does induce some postzygotic isolation, due to a disadvantage of rare mating types, a phenomenon discussed previously by Kawata and Yoshimura (2000). This is a rather weak effect, but in at least some models can drive reinforcement (Kirkpatrick and Servedio 1999; Otto et al. 2008). Whether reinforcement is driven by this or other causes of sufficiently low hybrid fitness, for instance as a by-product of sexual or natural selection in allopatry, on secondary contact postzygotic isolation could lead to the evolution of more prezygotic isolation. That selection for prezygotic isolation can then have an incidental effect on increased postzygotic isolation. This dynamic could lead to a positive-feedback loop, accelerating differentiation and isolation of the two populations.

However, the dynamics of selection on mating behavior in a hybrid zone are likely to be more complex than captured in modelling done to date. This is because the optimal mating behavior (and the strength of selection toward that optimum) depends on the relative density of different mating phenotypes (Wilson and Hedrick 1982; Otto et al. 2008), and therefore on the location of an individual in relation to the hybrid zone: On the forefront of the species range, it may be advantageous for individuals to have a wide mate acceptance curve that allows hybridization, because if conspecific mates are too difficult to find it may be favorable to mate with heterospecifics. Alleles conferring broad acceptance curves would then tend to spread between species, reducing prezygotic isolation. In areas with only one pure species, there is no selection on the mate acceptance curve, since there is no variation in mates (in the simplest case, at least). Only when both species are at reasonably high frequency (i.e., in the center of the contact zone) is there selection against hybridization when postzygotic isolation is sufficiently strong. We must also consider hybrids: for them, a wide mate acceptance curve may be favored, since their offspring may be more successful if they breed with a pure individual of one of the species. This effect also works against reinforcement. To understand the conditions under which reinforcement is predicted to be the net effect in hybrid zone, it will be necessary to model these dynamics using continuous space.

Do these results imply that sexual and social selection are unimportant early in speciation? Certainly not. While the emphasis in the literature on sexual and social selection has been on their contribution to prezygotic isolation, they can equally well produce postzygotic isolation (reviewed by Uy et al. 2018). Any process that drives evolution within two isolated populations can result in genetic incompatibilities that reduce fitness of hybrids. These can range from mild to severe, but the simulation results show that only a small reduction in hybrid fitness has a powerful effect in limiting the blending of two species. Sexual and social selection may be especially prone to lead to hybrids having characteristics that are not simply a blending of the two species but rather are surprising and outside the range of variation in the parental forms (i.e., “transgressive” traits; Rieseberg et al. 1999; Dittrich-Reed and Fitzpatrick 2013; Campagna et al. 2018). Finally, assortative mating itself can result in a form of selection against intermediate hybrids, due to rare-type disadvantage (e.g., Bridle et al. 2006). All of these forms of lower fitness of hybrids can be a result of sexual and social selection. While these findings suggest assortative mating, unless perfect or very nearly so, is ineffective in maintaining isolation of two species, they encourage researchers to examine the effects of sexual selection and sexual signals on postzygotic isolation. Ultimately, it is the fitness of hybrids that is crucial in determining the future of a hybrid zone.

### AUTHOR CONTRIBUTIONS

D.E.I. conceived of the study, designed and analyzed the simulations, and wrote the article.

## ACKNOWLEDGMENTS

I am grateful to Jessica Irwin and Trevor Price for extensive and insightful discussion; and the attendees of talks on this topic at the International Ornithological Congress and the Gordon Research Conference on Speciation for enthusiastic and encouraging feedback. I thank Dolph Schluter for advice on cline fitting. Insightful comments on an earlier version of the manuscript were provided by Daniel Bolnick, Jessica Irwin, Daniel Matute, Trevor Price, Ken Thompson, and two anonymous reviewers.

## DATA ARCHIVING

Following acceptance into a peer-reviewed journal, data and the HZAM code will be archived at Dryad. Supplemental material, including videos showing simulations, are available at: https://www.zoology.ubc.ca/~irwin/irwinlab/hzam-hybrid-zone-with-assortative-mating-modelling-august-2019-version/

## SUPPLEMENTAL FIGURES

**Figure S1.**
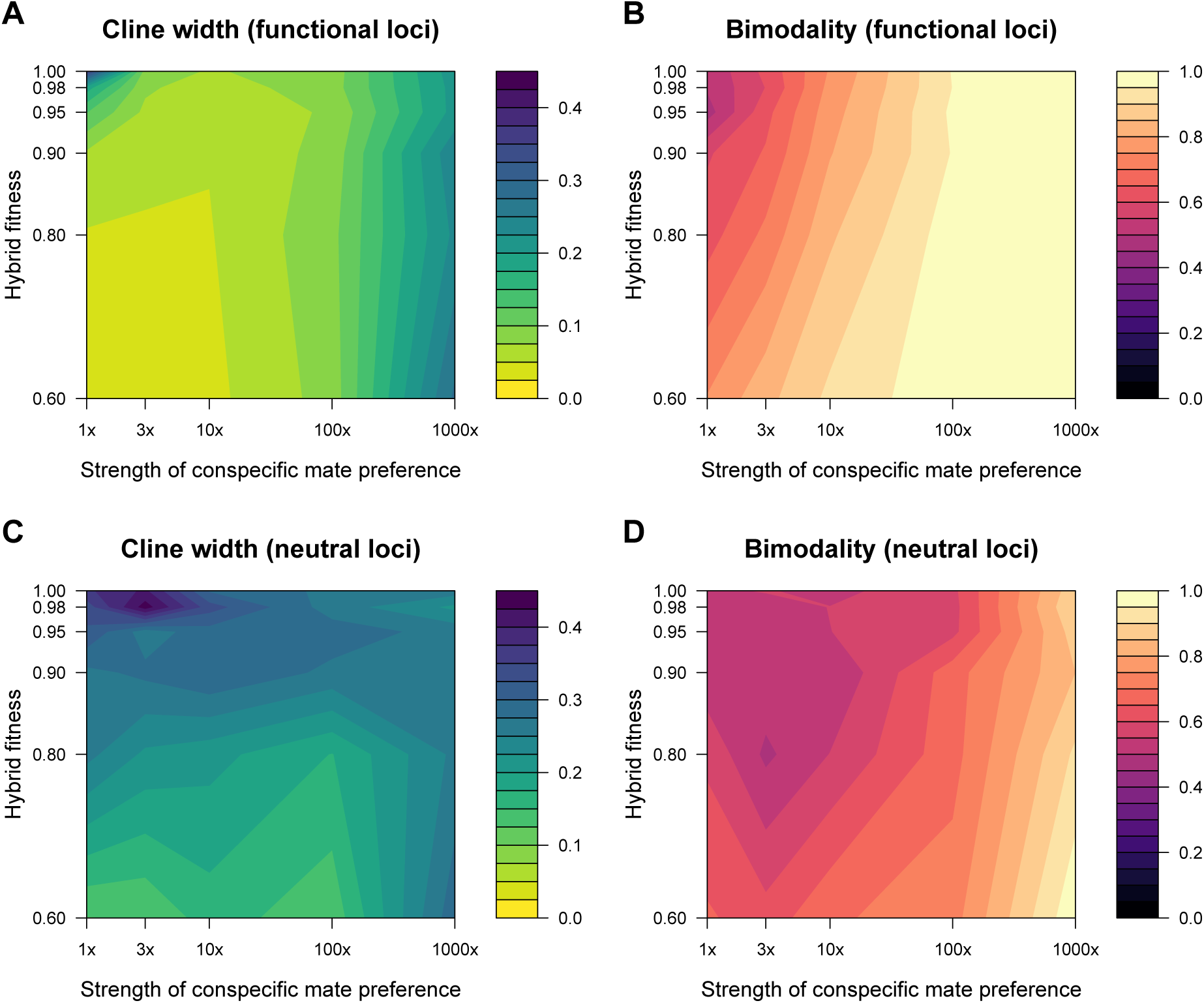
Summary of simulations run under identical conditions as those in Fig. 4, but with one functional locus and one neutral locus (instead of 3 of each). The cline in the functional locus narrows with increasing assortative mating more quickly in the 1-locus case than in the 3-loci case (see Figs. 4, 5).

**Figure S2.**
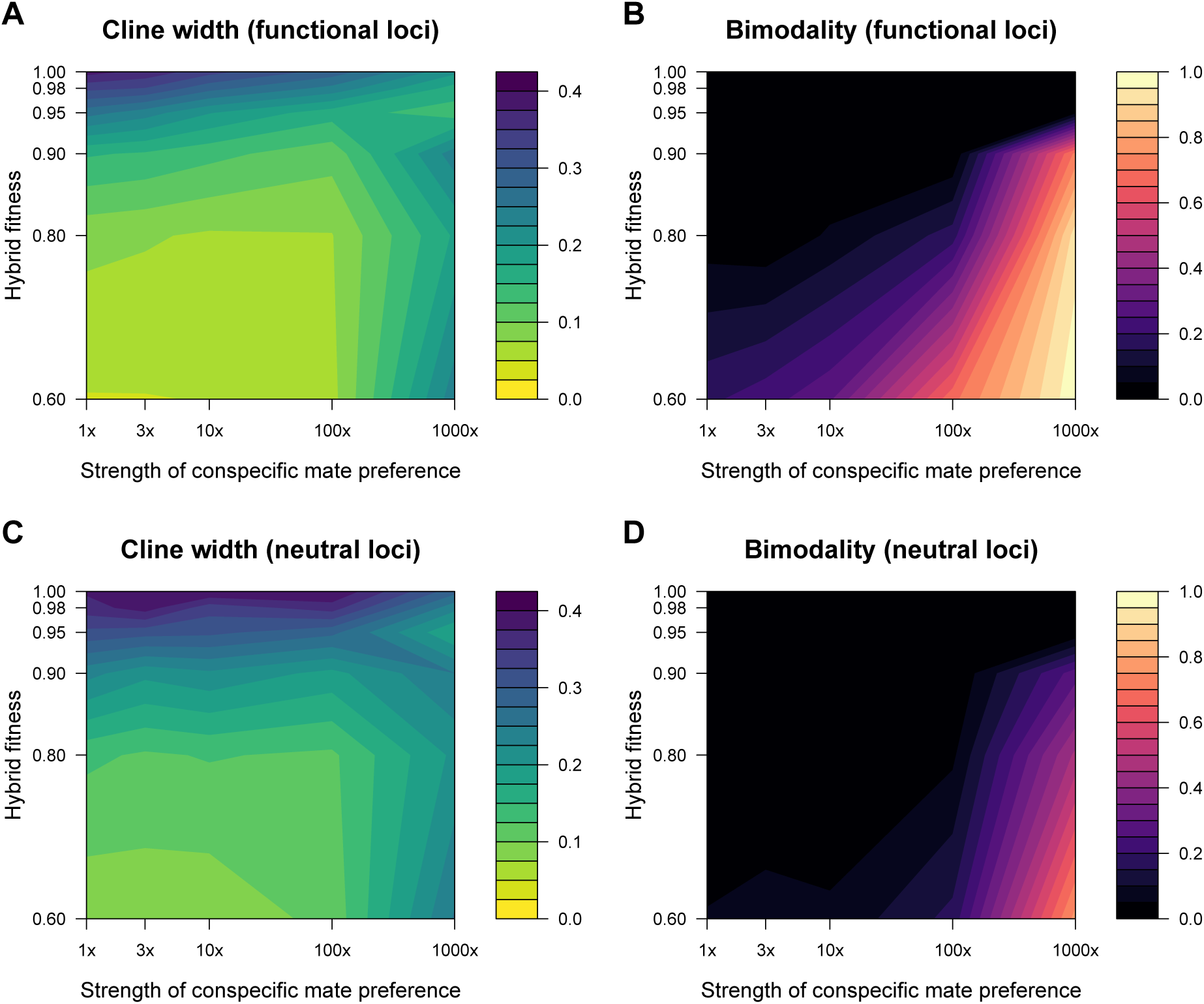
Summary of simulations run under identical conditions as those in Fig. 4, but with 9 functional loci and 9 neutral loci (instead of 3 of each). The width of the cline in the functional locus narrows with increasing assortative mating more slowly than in the 1-locus case and the 3-locus case (see Figs. 4, 5).

**Figure S3.**
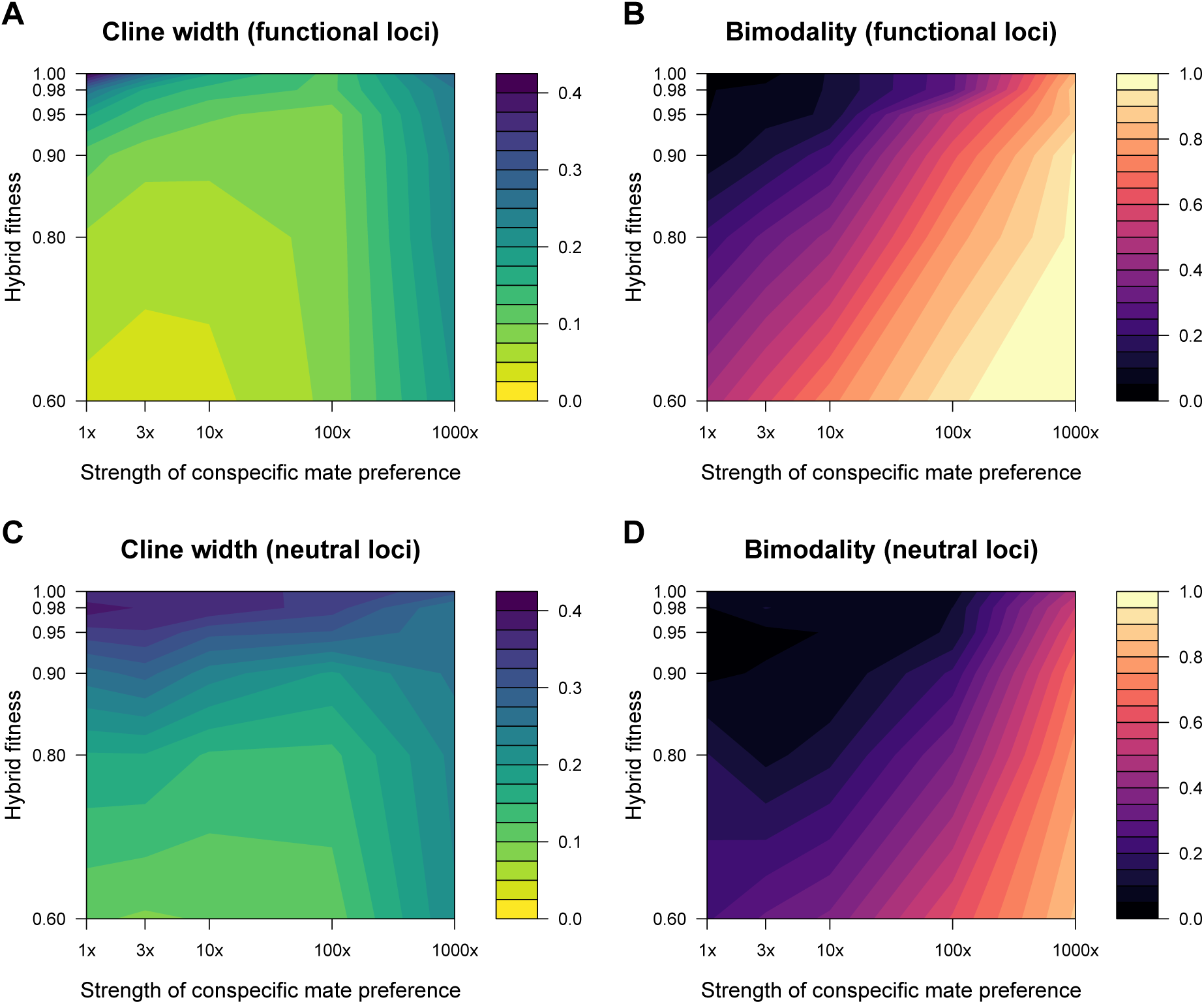
Summary of simulations run under identical conditions as those in Fig. 4, except the mating trait (both signal and preference) is encoded by loci that have dominant and recessive alleles (rather than codominant, in the basic model). At the start of simulations, species B is fixed for dominant alleles at each of the three loci contributing to the mating trait, and species A for recessive alleles. Results are similar to the codominant case, with the main difference being that neutral loci tend to flow more readily in the dominant case than the codominant case (compare panel C to Fig. 4C), likely due to the fact that F1 hybrids have a mating phenotype identical to one of the parental forms, providing a conduit for backcrossing and gene flow.

**Figure S4.**
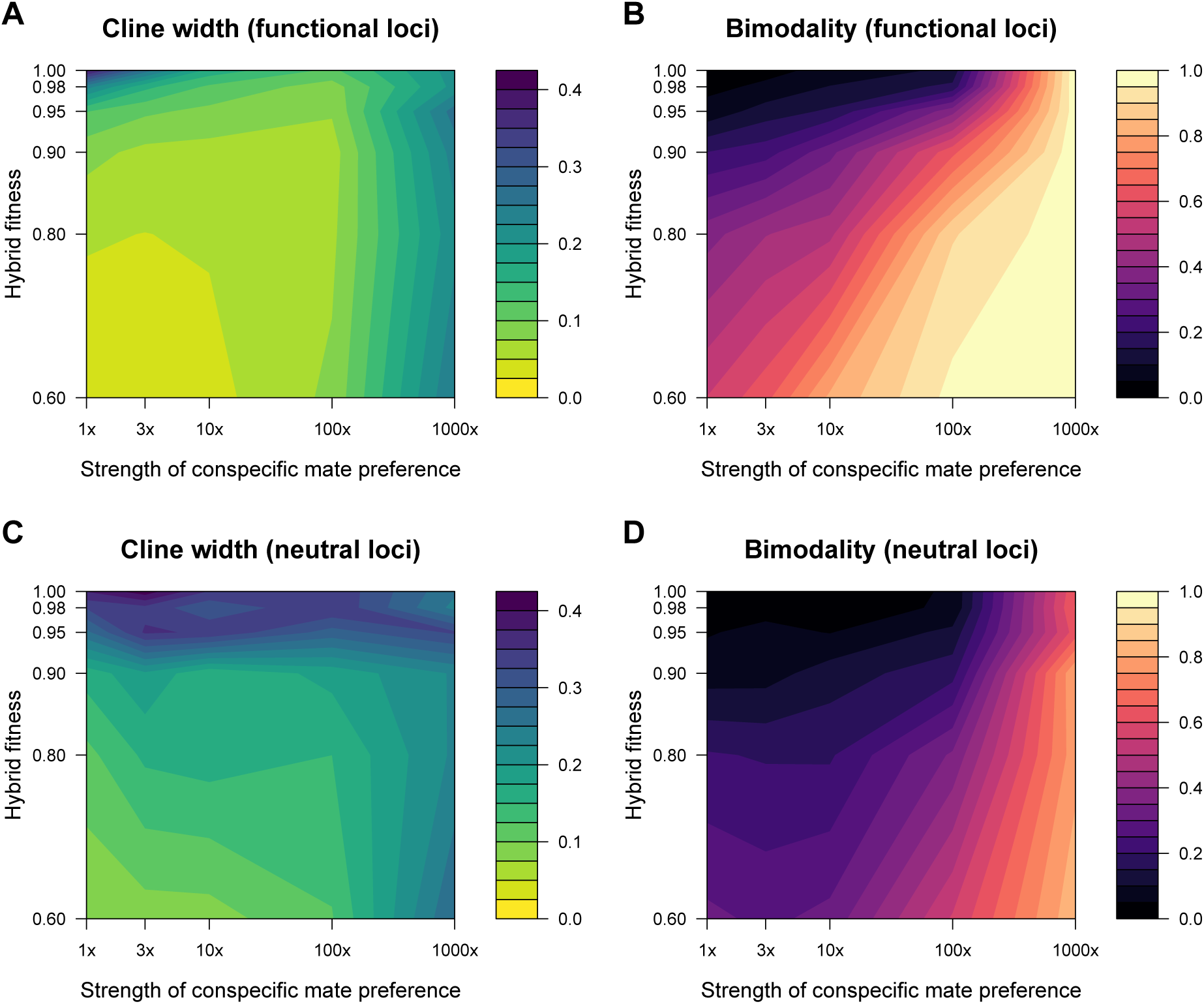
Summary of simulations run under identical conditions as those in Fig. 4, except that epistasis (interactions between loci) rather than simple underdominance determines survival probability. In this set of simulations, the epistasis parameter (*β*) is equal to 1.

**Figure S5.**
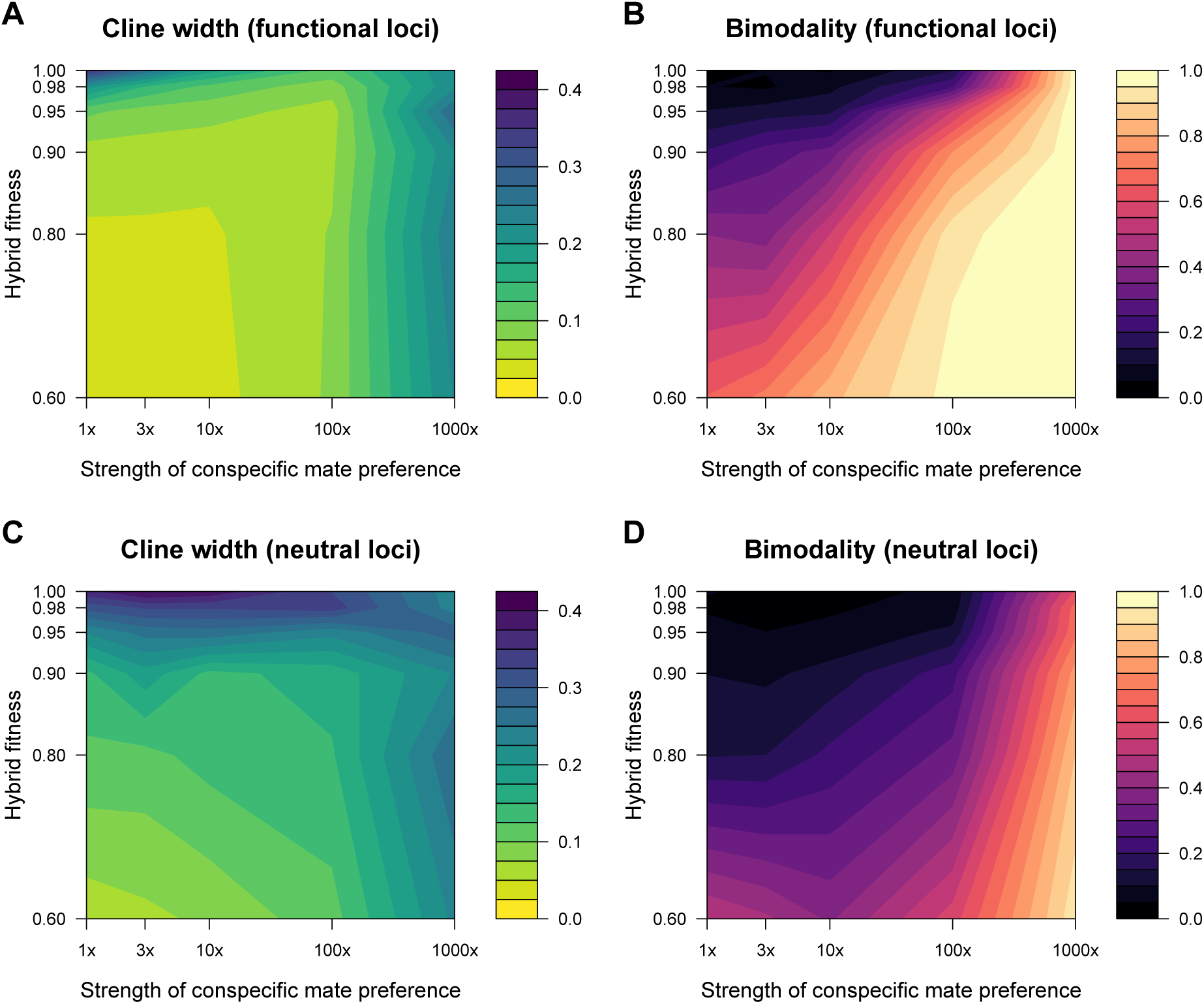
Summary of simulations run under identical conditions as those in Fig. 4, except that epistasis (interactions between loci) rather than simple underdominance determines survival probability. In this set of simulations, the epistasis parameter (*β*) is equal to 1/16 = 0.0625 (compare with Figs. S4 and S6, based on *β* of 1 and 16 respectively).

**Figure S6.**
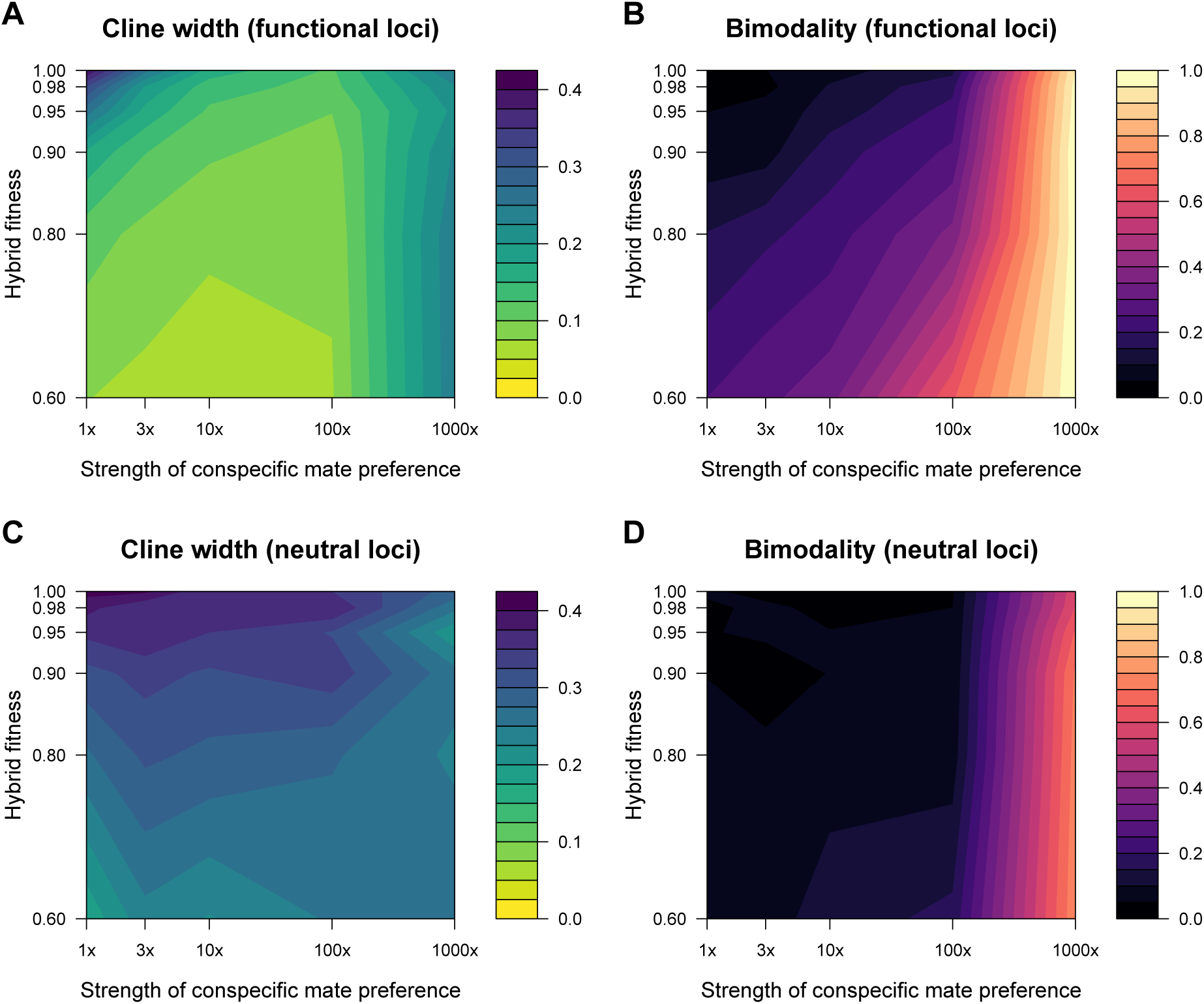
Summary of simulations run under identical conditions as those in Fig. 4, except that epistasis (interactions between loci) rather than simple underdominance determines survival probability. In this set of simulations, the epistasis parameter (*β*) is equal to 16 (compare with Figs. S4 and S5, based on *β* of 1 and 1/16 respectively).

**Figure S7.**
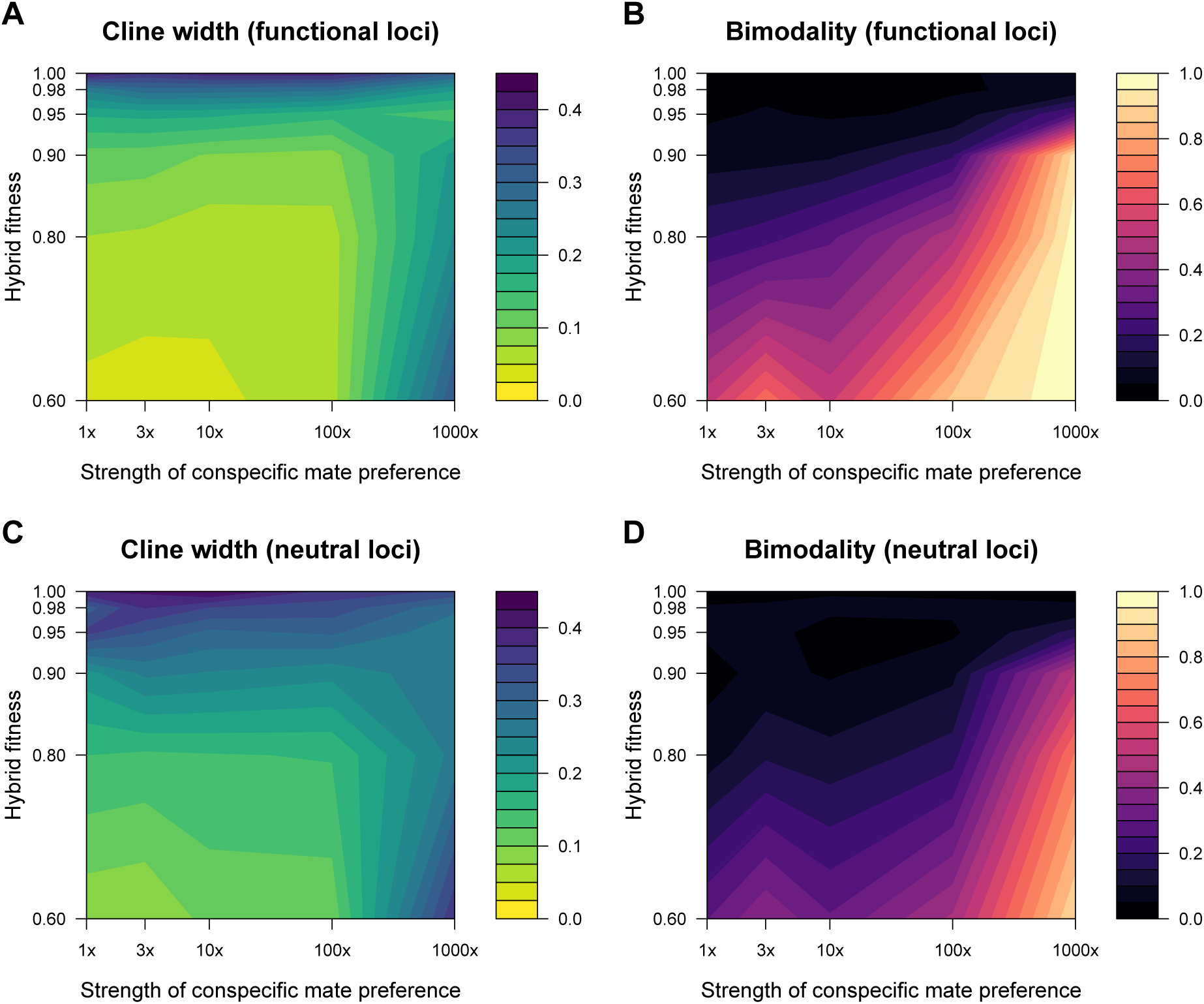
Summary of simulations run under identical conditions as those in Fig. 4, except that there are different loci (3 each) encoding the male trait and the female preference. In this case, when there is no reduced fitness of hybrids, there is virtually no impact of assortative mating on keeping a zone narrow and/or bimodal.

**Figure S8.**
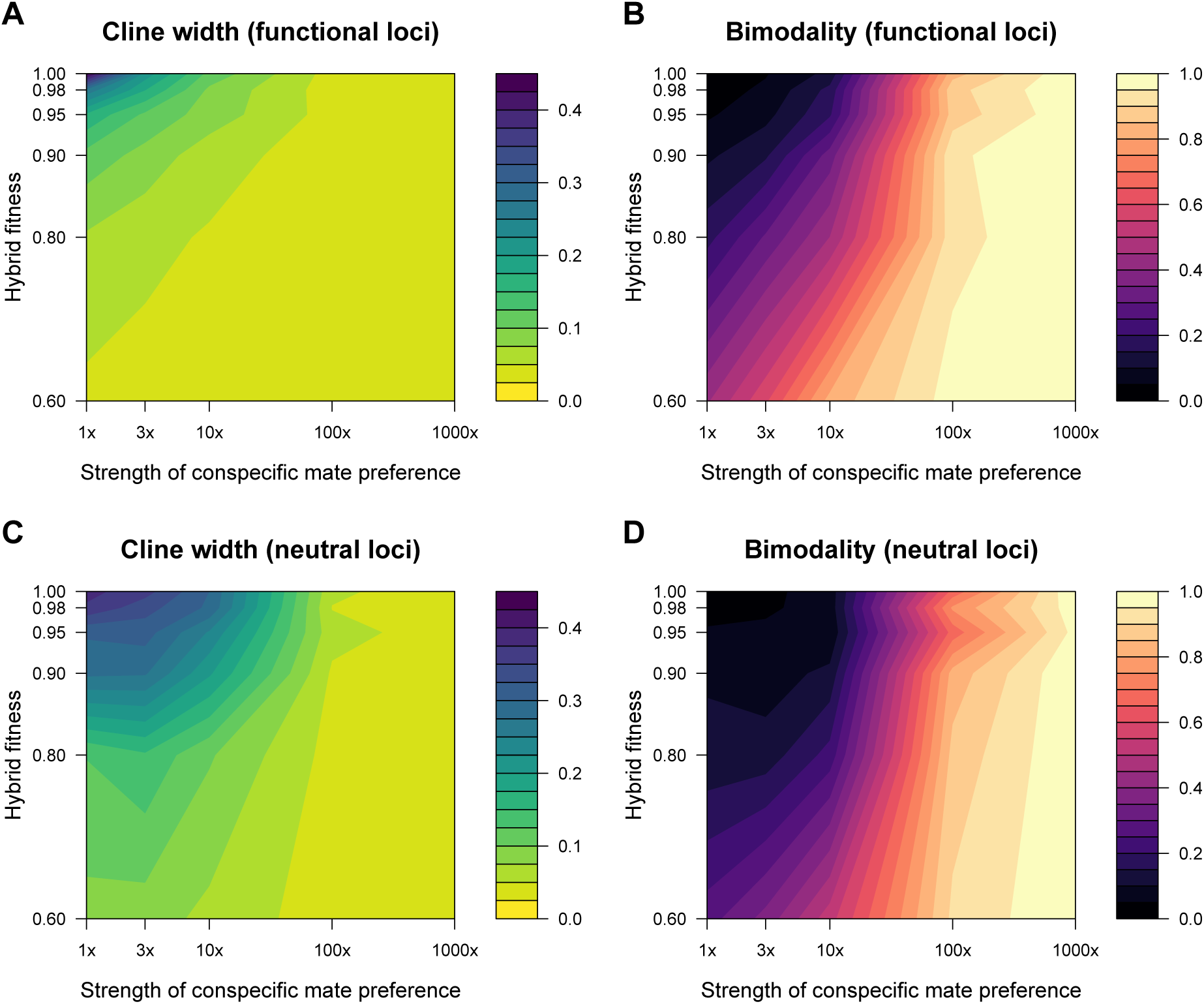
Summary of simulations run under identical conditions as those in Fig. 4, except that there is a mate search cost for females of 10% reduced fecundity per male rejected. This search cost causes mating preferences to have a much larger role in keeping a hybrid zone narrow and bimodal. The cost tends to be paid more by rare mating types—when hybrids are rare, they tend to pay this cost more than pure forms, hence it is a form of postzygotic isolation.

